# A minimal set of internal control genes for gene expression studies in head and neck squamous cell carcinoma

**DOI:** 10.1101/108381

**Authors:** Vinayak Palve, Manisha Pareek, Neeraja M Krishnan, Gangotri Siddappa, Amritha Suresh, Moni Abraham Kuriakose, Binay Panda

## Abstract

**Background:** Selection of the right reference gene(s) is crucial in the analysis and interpretation of gene expression data. In head and neck cancer, studies evaluating the efficacy of internal reference genes are rare. Here, we present data for a minimal set of candidates as internal control genes for gene expression studies in head and neck cancer.

**Methods:** We analyzed data from multiple sources (*in house* whole-genome gene expression microarrays, *n*=21; TCGA RNA-seq, *n*=42, and published gene expression studies in head and neck tumors from literature) to come up with a set of genes (discovery set) for their stable expression across tumor and normal tissues. We then performed independent validation of their expression using qPCR in 14 tumor:normal pairs. Genes in the discovery set were ranked using four different algorithms (BestKeeper, geNorm, NormFinder, and comparative delta Ct) and a web-based comparative tool, RefFinder, for their stability and variance in expression across tissues.

**Results:** Our analyses resulted in 18 genes (discovery set) that had lowest variance and high level of expression across tumor and normal samples. Independent experimental validation and analyses with multiple tools resulted in top ranked five genes (*RPL30, RPL27, PSMC5, OAZ1* and *MTCH1*) out of which, *RPL30* (60S ribosomal protein L30) and *RPL27* (60S ribosomal protein L27), performed best and were abundantly expressed across tumor and normal tissues.

**Conclusions:** *RPL30* and *RPL27* are stably expressed in HNSCC and should be used as internal control genes in gene expression in head and neck tumors studies.

## Background

Internal control genes or housekeeping genes are important in obtaining reliable and reproducible data from gene expression studies ^1^. Internal control genes should be abundantly and uniformly expressed across tumor and normal tissues and across different experimental conditions ^2^. Head and neck squamous cell carcinoma (HNSCC) is the sixth most common cancer worldwide with an incidence of 550,000 cases annually ^3^. Although there are reports on reference internal control genes in head and neck tumor gene expression studies ^4^, studies based on discovery from high-throughput data like microarrays and RNA-seq, the confirmation of their expression using qPCR across tumor and normal samples are absent. Past studies argue the lack of uniformity of expression on internal control genes based on experimental conditions ^5 6^. Therefore, it is crucial that the expression of internal control genes remains unaltered across temporal, spatial and experimental conditions. Additionally, internal control genes need to take into account genes expressed both at high and low level of expression. Availability of high-throughput gene expression data from different cohorts and large consortia like TCGA warrants revisiting the validity of widely used genes like *ACTB*, *TUBB* and *GAPDH*.

In the present study, we analyzed HNSCC gene expression data from three sources; *in house* microarray data, TCGA RNA-seq data and data from literature to come up with a set of genes (discovery set) that are stably and robustly expressed with least variance across tumor: normal pairs. We subsequently validated the expression of the discovery set in tumor: normal pairs (*n*=14) independently using qPCR. Finally we obtained a list of two genes as a minimal set by comparing and ranking their expression during validation.

## Methods

### Microarray data processing

The gene expression profiling using Illumina HumanHT-12 v4 expression BeadChips (Illumina, San Diego, CA), raw data collection and data processing using the R package lumi ^7^ and individual batch-correction using ComBat ^8^ is described previously ^9 10^. The raw signal intensities from arrays were exported from GenomeStudio for pre-processing and analyzed using R. Gene-wise expression intensities for tumor and matched control samples from GenomeStudio were transformed and normalized using VST (Variance Stabilizing Transformation) and LOESS methods, respectively, using lumi ^7^ and top genes with least across-sample variance in expression were selected. Genes with fold change between 0.95-1.05 (tumor/normal) and with standard deviation < 0.05 for the fold changes were selected further.

### RNA-seq data processing

RSEM-processed RNA-Seq gene expression values (TCGA pipeline, Level 3) were downloaded from the old TCGA repository (https://genome-cancer.ucsc.edu/proj/site/hgHeatmap/). The TPM values for total 42 matched tumor-normal pairs for all the genes were extracted from the Level 3 files. Further, genes that showed zero or *NA* values in any of these samples were eliminated, and the log fold change values between the respective tumor and normal samples calculated by taking a log transformation of the ratios between expression values for tumor and normal. We filtered expression data that fulfilled three criterial for all samples; expressed at >= 3 TPM, tumor/normal ratio: 0.9-1.1 and standard deviation across all the tissues < 0.5. Finally genes were short-listed based on their levels of expression across tumor: normal pairs, low standard deviation across samples, are well annotated and with some level of biology known.

### Discovery set

In addition to the microarray and RNA-seq gene expression data, the most commonly used internal control genes form the HNSCC literature (*ACTB*, *TUBB* and *GAPDH*) were also considered. Genes that satisfied all the above criteria plus the commonly used genes in the literature resulted in a list of 18 genes, discovery set (**Table 1
**) that was used for validation in an independent set of tumor: normal pairs.

### Patient samples

In the discovery and validation stage, HNSCC tumor: adjacent normal paired samples (*n*=63 and *n*=14 respectively), were used. Previously published whole-genome microarray data ^9 10^, RNA-seq data from TCGA ^11^ and data from individual studies in the literature were used in the discovery stage. For the validation set, 10 HNSCC tumor: normal pairs were from a completely independent cohort (which were not a part of discovery cohort) and 4 were used from the *in house* discovery cohort. For the independent validation cohort, patients were accumulated voluntarily after obtaining informed consents from each patient and as per the guidelines from the institutional ethics committee. All the tissues were frozen immediately in liquid nitrogen and stored at −80°C until further use. Only those tumors with squamous cell carcinoma, with at least 70% tumor cells and confirmed diagnosis were included in the current study. Patients underwent staging according to AJCC criteria, and curative intent treatment as per NCCN guidelines involving surgery with or without post-operative adjuvant radiation or chemo-radiation at the Mazumdar Shaw Cancer Centre were accrued for the validation study.

### RNA extraction and cDNA synthesis

The total RNA was extracted from 25mg of tissues using the RNeasy mini kit (Qiagen, cat:74104) with on-column digestion of DNA using RNase free DNase set (Qiagen, cat:79254) as per manufacturer’s instructions. The RNA quality was checked using gel electrophoresis and Agilent Bioanalyzer. Five hundred nanogram of total RNA was subjected to cDNA synthesis using Takara’s Prime Script first strand cDNA synthesis kit (cat: 6110A).

### Quantitative real-time PCR (qPCR)

Quantitiatve PCR was carried out using KAPA Biosystem’s SYBR Fast qPCR universal master mix (cat: KK4601). The primer sequences for all the 18 candidate reference genes and the amplification conditions are mentioned in Table 1A, which were either designed for this study, were chosen from the literature ^12^ or from online resources (https://primerdepot.nci.nih.gov/ and https://pga.mgh.harvard.edu/primerbank/). All amplification reactions were carried out in triplicates, using three negative cotrols: no template control (NTC) with nuclease free water (Ambion, cat: AM9932), no amplification control (NAC) and no primer control (NPC) in each amplification plate.

### Statistical analysis

For stability comparison of the internal candidate reference genes obtained using the discovery set, we analyzed the validation data using four most commonly used algorithms, Genorm ^13^, Normfinder ^14^, Bestkeeper ^15^ and Comparative Ct ^16^ were used and genes were compared and ranked using RefFinder ^17^. Graphpad prism software version 5 was used to analyze the data and plot the graphs.

## Results and Discussion

The schema for selecting genes for discovery set is depicted in Figure 1. After analyzing data from all sources, 18 genes were selected that had least variance across all the tumor and normal samples, were well annotated and had some biology known (Table 1A and Supplementary Table 1). Genes in the validation tumor: normal pairs, we produced standard curves reflecting the linear regression (R^2^ >0.9) and amplification efficiency (Supplementary Figure 1). Primers for all the genes showed specific amplifications as shown by the dissociation curves in Supplementary Figure 2. Data from qPCR validation of genes (Figure 2A) showed variable levels of expression for the 18 genes in different validation samples. As Figure 2B indicates, results from the four different algorithms (Genorm, Normfinder, Bestkeeper & Comparative Ct) and a comprehensive RefFinder tool showed expression of 5 genes (*RPL30*, *RPL27*, *PSMC5*, *OAZ1* and *MTCH1*) were highly consistent across all the samples among the discovery panel. Further, we ranked the genes using the above tools (Figure 2B) and 11 genes (*RPL30*, *RPL27*, *PSMC5*, *OAZ1*, *MTCH1*, *TSPAN21*, *DARS*, *MKRN1*, *RPS13*, *RPL5* and *RPL37A*) ranked on the top depending on the tool used. Using the tool RefFinder, that takes into account all other algorithms and performs its own calculation to come up with a rank, we got 5 candidate reference genes (*RPL30*, *RPL27*, *PSMC5*, *OAZ1*, *MTCH1*) as most stable in terms of their expression. Two genes out this list, *RPL27* and *RPL30*, fulfilled all the 3 criteria for an ideal internal control set, least variation in their expression across samples, high-level of expression in both tumors and normal and ranked top by the algorithm.

**Figure 1:**
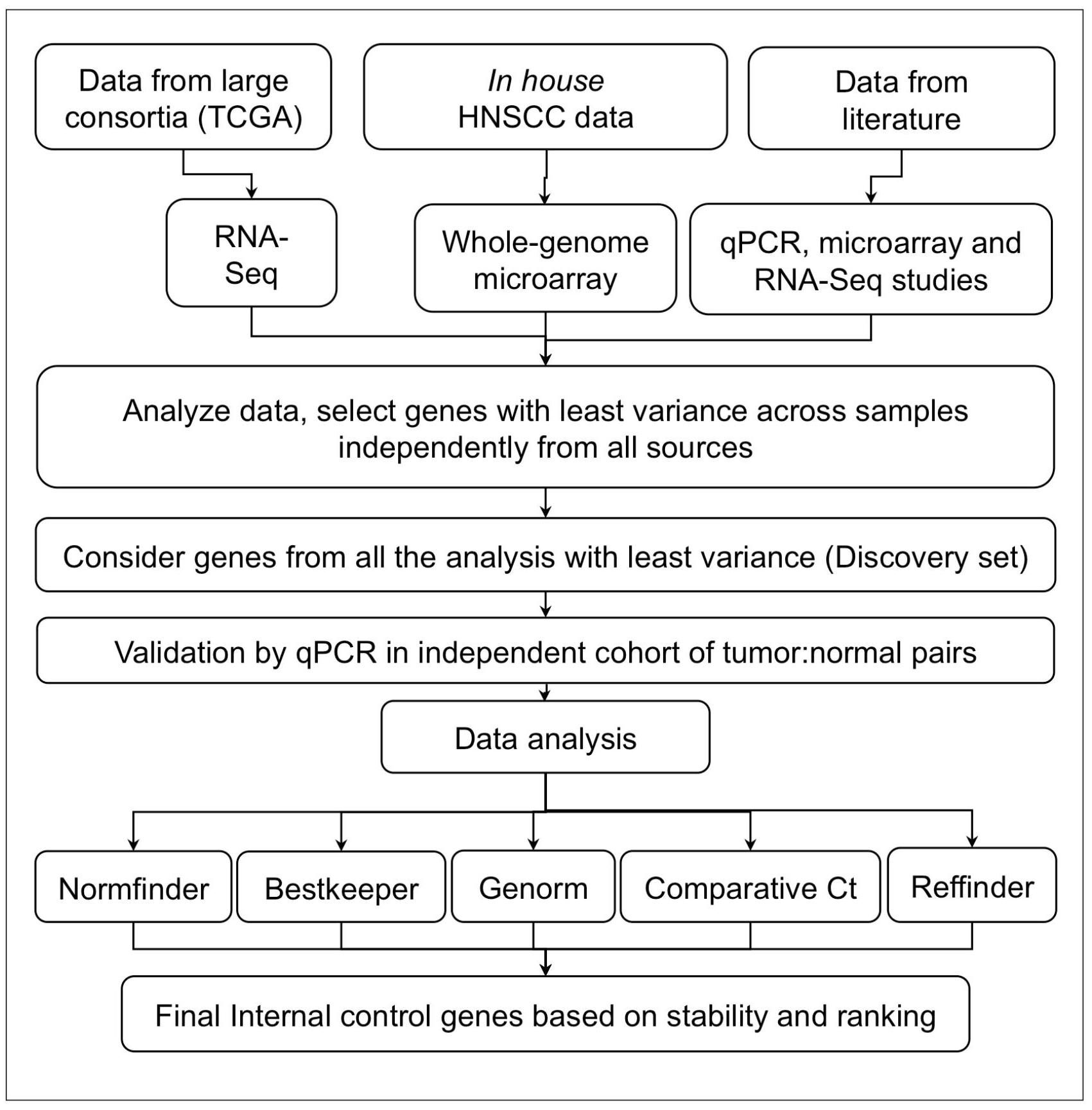
Schema indicating the selection criteria and validation of candidate internal control genes.

**Figure 2:**
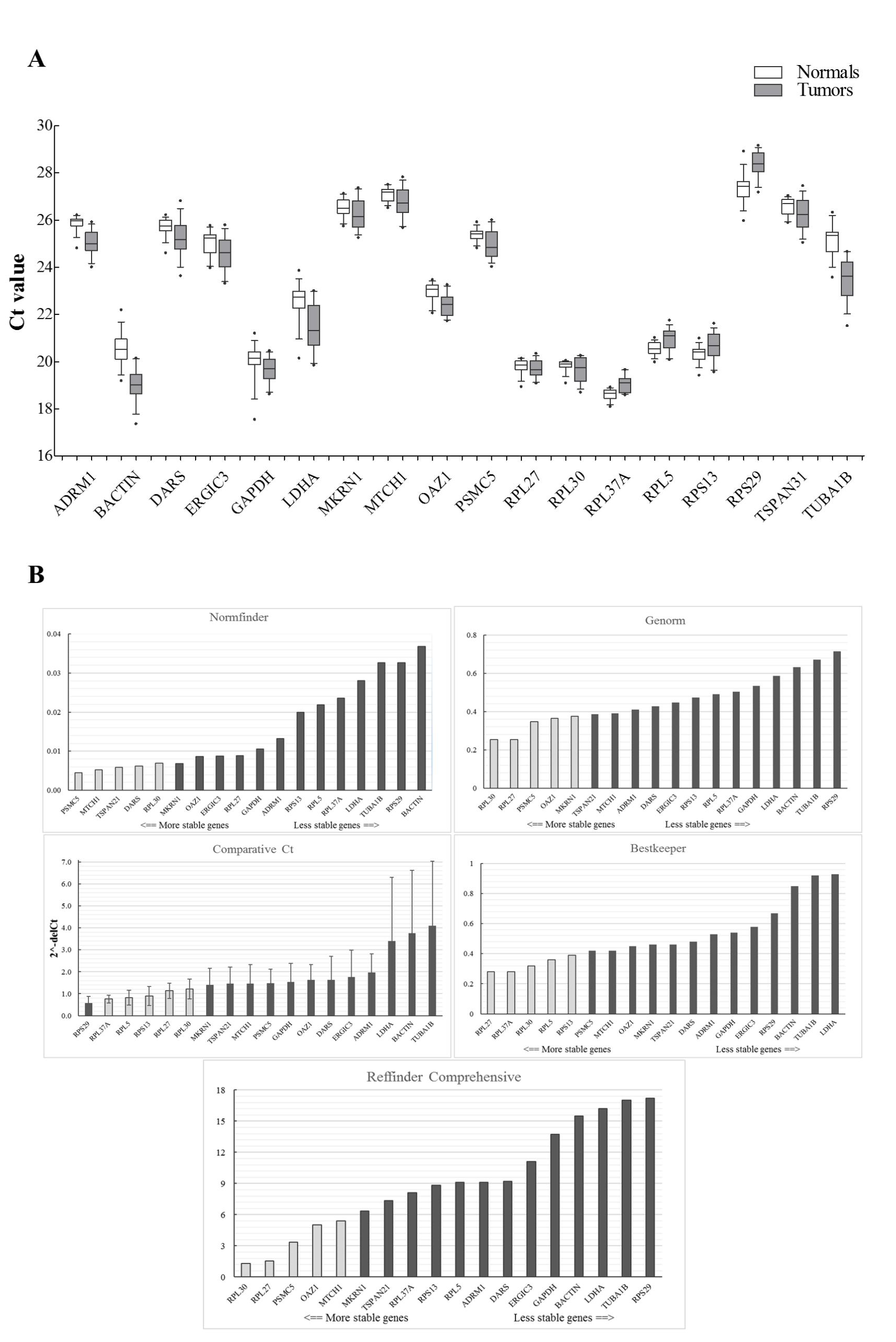
Expression of genes in the discovery set across tumors and normal (A).The Ct values were generated from qPCR experiments and plotted as box plots with median values as lines and boxes indicating 10-90 percentile values and the whiskers as the maximum and minimum values. Ranking of 18 genes using different tools (B). Y-axis for different tools represents; stability value (Genorm, Normfinder), Pearson correlation coefficient (Bestkeeper), 2-^∆∆Ct^ (Comparative Ct method) and geometric ranking values (RefFindser). The columns in light grey represents stable genes across tumors and normal samples.

**Table 1:**
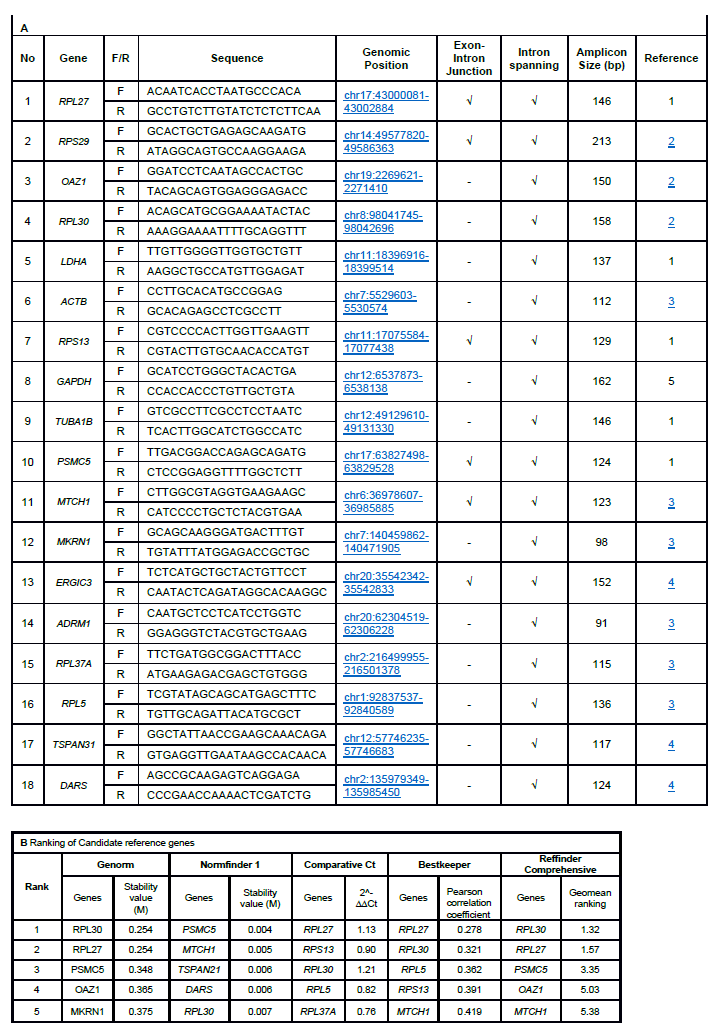
Primers used in the current study (A). The numbers in the Reference column indicate-1: primers designed for the study; 2: primers from Hendrik de Jonge et al., 2007; 3: primers from Primerdepot (https://primerdepot.nci.nih.gov/); 4: primers from primer bank (https://pga.mgh.harvard.edu/primerbank/); 5: primers from Campos et al, 2009;. The top ranked genes and their ranks as obtained from using various software tools (B).

The quantitative PCR is a highly sensitive and widely used method employed for studying gene expression, especially validation of gene expression data. Several factors may affect the qPCR data including, RNA input volume, cDNA synthesis efficiency, pipetting volumes and accuracy, and primer amplification efficiencies ^18^. Use of internal control genes overcomes these shortcomings and helps interpret data reliably and reproducibly. Although past studies are heavily reliant on the use of *ACTB*, *TUBB* and *GAPDH* to normalizes the qPCR data, there are evidence that the expression of these genes vary greatly and are unsuitable to be used as internal control genes ^19#22^. Additionally, expression of internal control genes may vary based on experimental conditions and tumor type. Therefore, it is important to have a set of genes that can be used across tumors and normal samples in a cancer type. In HNSCC, previous attempts have been made to come up with a set of internal control genes using qPCR data ^4^. However, it is important that the discovery is made from a transcriptome-wide study that takes into account data from RNA-seq and microarrays. This is especially important as we have data on HNSCC from large consortia, like TCGA, and other high-throughput studies available. Keeping this in mind, we analyzed data from multiple high-throughput platforms (RNA-seq data from TCGA and *in house* genome-wide microarray data), multiple geographical cohorts and previously published data using different platforms (qPCR, microarray and others). We selected 18 genes (discovery set), the expression of which were stable and least variable across the samples (tumor: normal pairs) and were essential and code for structural proteins (ribosomal, cell structure and integrity), essential functions like glycolysis and translation (Supplementary Table 1). To the best of our knowledge, this study represents the first report defining a minimal set of robust candidate reference genes for gene expression studies in HNSCC that utilizes discovery from high-throughput studies followed by validation in an independent cohort. In our validation study, we found *RPL30*, *RPL27*, *PSMC5*, *OAZ1* and *MTCH1* as the top 5 ranked genes with stable expression across all the samples. Previous studies with head and neck cancer patients also found *RPL27* to be one the most stable candidate internal control gene^4^. Meta-analysis with 13,629 array samples derived from primary patient materials, cell lines, diseased as well as normal tissues, and with varying experimental conditions) have featured *RPL27* and *RPL30* in their list of top 5 housekeeping genes ^5^. Additionally, previous findings on non HNSCC tumors also had *RPL27*, *RPL30* and *MTCH1* in the list of top internal control genes ^23^. Although, *PSMC5*, *OAZ1* and *MTCH1* were stably expressed and ranked after *RPL30* and *RPL27* by RefFinder, they were expressed at a lower level than *RPL30* and *RPL27*. *RPL30*, *RPL27*, however, were stably and abundantly expressed across all the tumor and normal samples with least variance (Figure 2A). Therefore, we term them as a minimal internal control gene set for HNSCC studies. Although both *RPL30* and *RPL27* were equally good, in resource-poor settings and/or where sample amounts are limiting, any one of them can equally serve as an internal control gene. Although our study reveals two robust internal control genes post discovery from large high-throughput studies followed by independent validation, it suffers from some shortcomings. First, several factors may influence our selection of discovery set, difference of gene expression due to inter-platform and assay variability along with a smaller sample size. A larger sample size used by a single platform and kit (for example, total transcriptome RNA-seq using a single provider’s platform and assay version) may solve this in the future. Second, qPCR experiments depend on the primer design, locations of primers, amplification of transcript variants (if any) and all or any of these could have influenced our results from the validation step. Third, our validation sample size was small and there is a possibility that we may get a larger variability if a larger validation set is used. Fourth, our results could have been biased due to the type of algorithm/software used to analyze the qPCR data. In our study, we relied on the RefFinder tool for the ranking of the genes as it is comprehensive and provided a gene list that mostly overlapped with the list provided by the other 4 tools. With the exception of NormFinder, *RPL30* and *RPL27* were in the top 5 genes by all the tools. Like previously reported ^25^, a difference in the ranking of genes by different tools may result due to lack of consideration of certain parameters (like qPCR amplification efficiency) by certain tools (like RefFinder).

## Conclusions

Commonly used internal control genes like *ACTB*, *TUBB*, *GAPDH* are not suitable reference genes for HNSCC gene expression studies. *RPL30* and/or *RPL27* represent a set of robust and stably expressed genes across tumor and normal samples and should be used as internal control/housekeeping genes in HNSCC gene expression studies.

## List of abbreviations

HNSCC: head and neck squamous cell carcinoma

## Authors' contributions

VP: Designed and performed experiments, analyzed data, wrote the manuscript; MP: performed experiments; NMK: analyzed data; GS, AS, MAK: procured patient materials; and BP: Conceived the study, provided overall study guidance, wrote the manuscript.

## Acknowledgement

The authors thank Janani Hariharan in helping with the RNA-seq data analyses. Research presented in this article is funded by Department of Electronics and Information Technology, Government of India (Ref No: 18(4)/2010-E-Infra., 31-03-2010) and Department of IT, BT and ST, Government of Karnataka, India (Ref No: 3451-00-090-2-22).

---

**Supplementary Figure S1:**
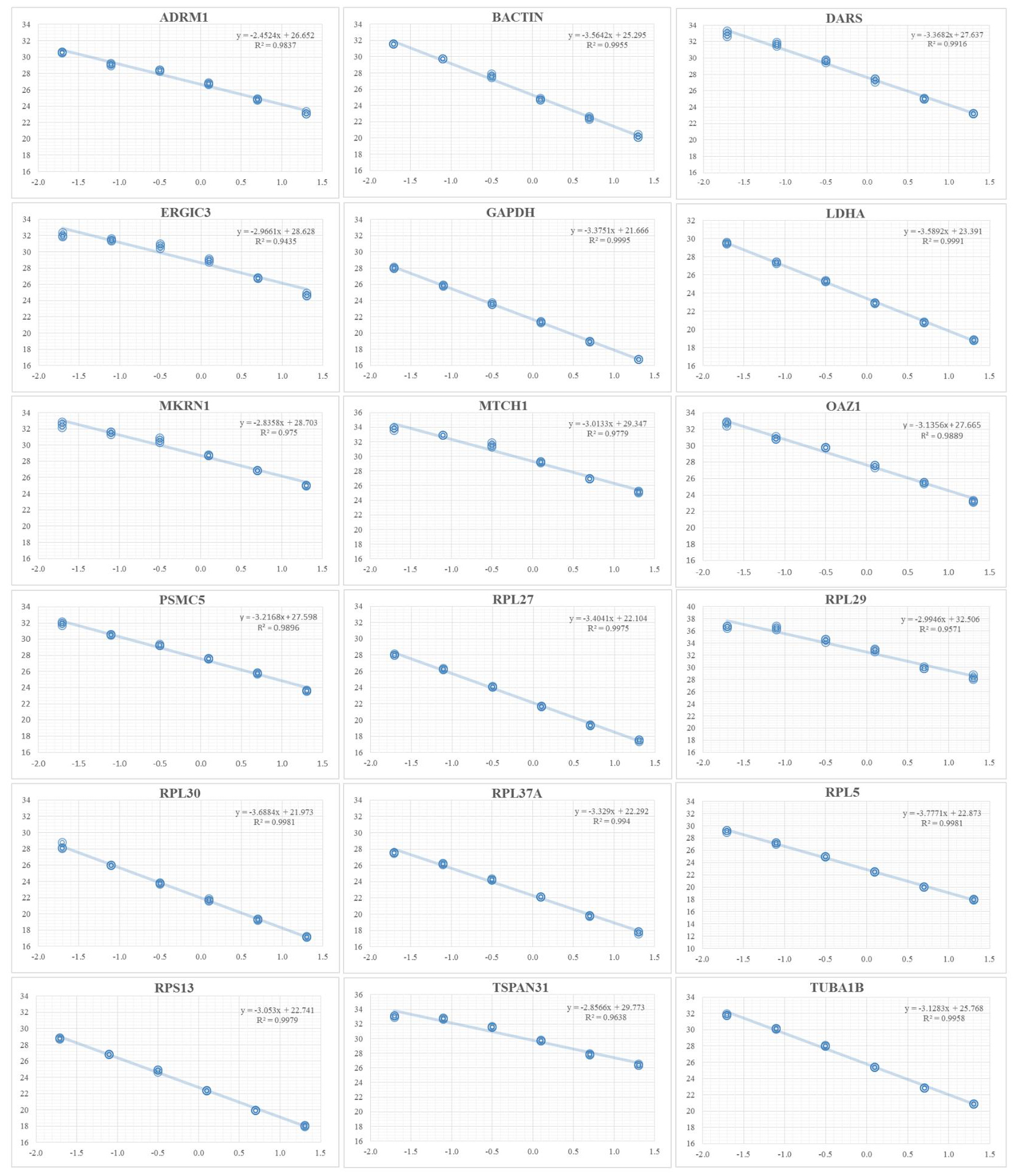
Standard curves of 18 candidate housekeeping genes.The qPCR was performed using 3 times dilution of the template. R2 indicates the correlation coefficient and amplification efficiencies on the basis of the slopes.

**Supplementary Figure S2:**
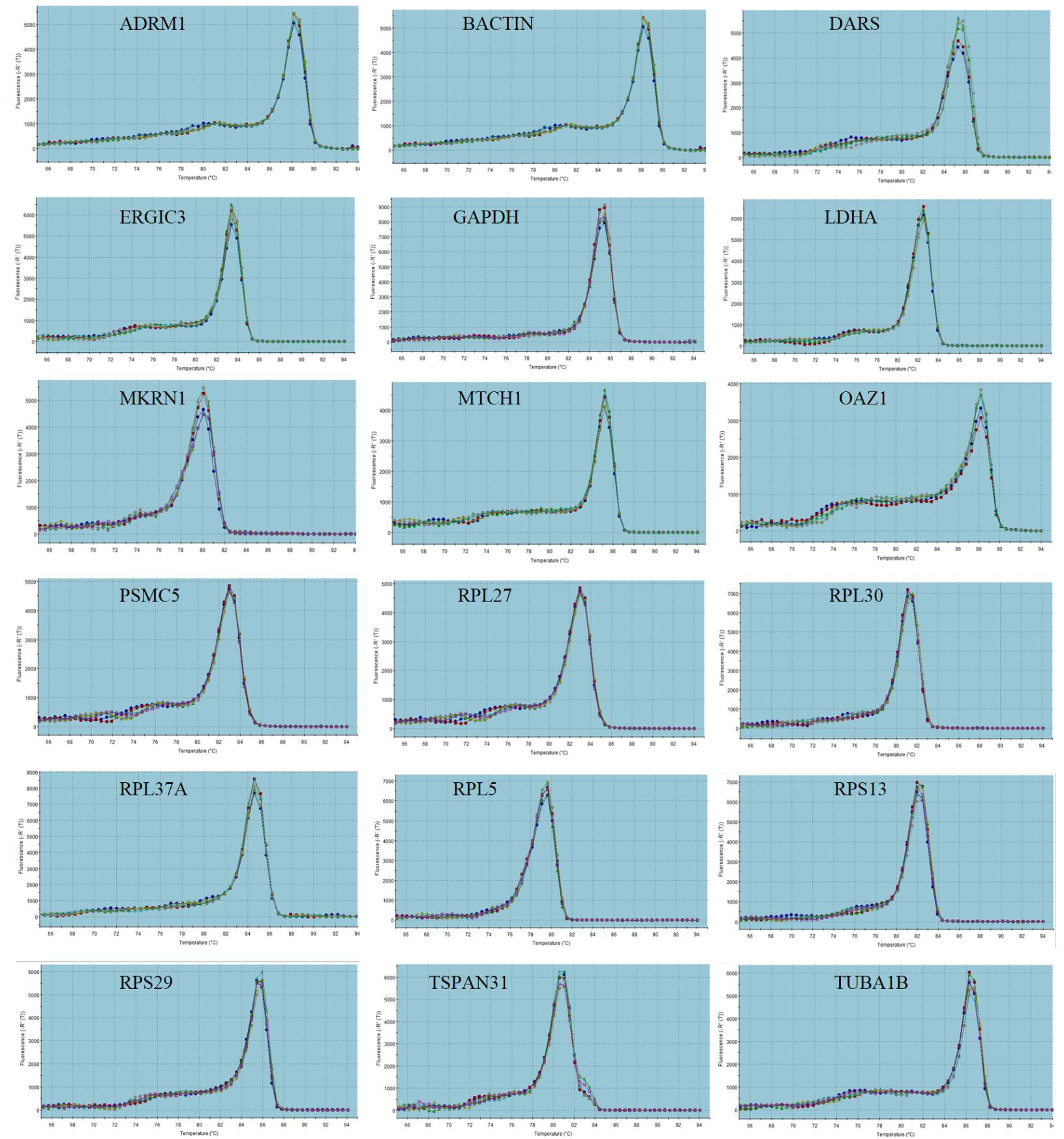
Dissociation curves for the 18 candidate housekeepinggenes. The single peak of dissociation curve in qPCR performed for all the genes indicates the specific amplification.

**Table S1.** Genes selected in the discovery set and their function.

| Supplementary Table 1: Genes selected in the discovery set and their function. |  |  |
| --- | --- | --- |
| Gene Name | Gene Functions | Source |
| RPL30 | This gene encodes a ribosomal protein that is a component of the 60S subunit. The protein belongs to the L30E family of ribosomal proteins. It is located in the cytoplasm. This gene is co-transcribed with the U72 small nucleolar RNA gene, which is located in its fourth intron. As is typical for genes encoding ribosomal proteins, there are multiple processed pseudogenes of this gene dispersed through the genome. | https://www.ncbi.nlm.nih.gov/gene/6156 |
| RPL27 | This gene encodes a ribosomal protein that is a component of the 60S subunit. The protein belongs to the L27E family of ribosomal proteins. It is located in the cytoplasm. As is typical for genes encoding ribosomal proteins, there are multiple processed pseudogenes of this gene dispersed through the genome. RPL27 (Ribosomal Protein L27) is a Protein Coding gene. Among its related pathways are Viral mRNA Translation and Infectious disease. GO annotations related to this gene include poly(A) RNA binding and structural constituent of ribosome | https://www.ncbi.nlm.nih.gov/gene/6155 |
| RPS29 | This gene encodes a ribosomal protein that is a component of the 40S subunit and a member of the S14P family of ribosomal proteins. The protein, which contains a C2-C2 zinc finger-like domain that can bind to zinc, can enhance the tumor suppressor activity of Ras-related protein 1A (KREV1). It is located in the cytoplasm. Variable expression of this gene in colorectal cancers compared to adjacent normal tissues has been observed, although no correlation between the level of expression and the severity of the disease has been found. As is typical for genes encoding ribosomal proteins, there are multiple processed pseudogenes of this gene dispersed through the genome. Alternatively spliced transcript variants encoding different isoforms have been found for this gene. RPS29 (Ribosomal Protein S29) is a Protein Coding gene. Diseases associated with RPS29 include diamond-blackfan anemia 13 and diamond-blackfan anemia. Among its related pathways are Viral mRNA Translation and Activation of the mRNA upon binding of the cap-binding complex and eIFs, and subsequent binding to 43S. GO annotations related to this gene include structural constituent of ribosome. | https://www.ncbi.nlm.nih.gov/gene/6235 |
| OAZ1 | The protein encoded by this gene belongs to the ornithine decarboxylase antizyme family, which plays a role in cell growth and proliferation by regulating intracellular polyamine levels. Expression of antizymes requires +1 ribosomal frameshifting, which is enhanced by high levels of polyamines. Antizymes in turn bind to and inhibit ornithine decarboxylase (ODC), the key enzyme in polyamine biosynthesis; thus, completing the auto-regulatory circuit. This gene encodes antizyme 1, the first member of the antizyme family, that has broad tissue distribution, and negatively regulates intracellular polyamine levels by binding to and targeting ODC for degradation, as well as inhibiting polyamine uptake. Antizyme 1 mRNA contains two potential in-frame AUGs; and studies in rat suggest that alternative use of the two translation initiation sites results in N-terminally distinct protein isoforms with different subcellular localization. Alternatively spliced transcript variants have also been noted for this gene. | https://www.ncbi.nlm.nih.gov/gene/4946 |
| LDHA | The protein encoded by this gene catalyzes the conversion of L-lactate and NAD to pyruvate and NADH in the final step of anaerobic glycolysis. The protein is found predominantly in muscle tissue and belongs to the lactate dehydrogenase family. Mutations in this gene have been linked to exertional myoglobinuria. Multiple transcript variants encoding different isoforms have been found for this gene. The human genome contains several non-transcribed pseudogenes of this gene. | https://www.ncbi.nlm.nih.gov/gene/3939 |
| ACTB | his gene encodes one of six different actin proteins. Actins are highly conserved proteins that are involved in cell motility, structure, and integrity. This actin is a major constituent of the contractile apparatus and one of the two nonmuscle cytoskeletal actins. Actins are highly conserved proteins that are involved in various types of cell motility and are ubiquitously expressed in all eukaryotic cells. | https://www.ncbi.nlm.nih.gov/gene/60 |
| TUBB (TUB1 | This gene encodes a beta tubulin protein. This protein forms a dimer with alpha tubulin and acts as a structural component of microtubules. Mutations in this gene cause cortical dysplasia, complex, with other brain malformations 6. Alternative splicing results in multiple splice variants. There are multiple pseudogenes for this gene on chromosomes 1, 6, 7, 8, 9, and 13. Tubulin is the major constituent of microtubules. It binds two moles of GTP, one at an exchangeable site on the beta chain and one at a non-exchangeable site on the alpha chain. Tubulin is the major constituent of microtubules. It binds two moles of GTP, one at an exchangeable site on the beta chain and one at a non-exchangeable site on the alpha chain. | https://www.ncbi.nlm.nih.gov/gene/203068,<br>https://www.ncbi.nlm.nih.gov/gene/10376 |
| GAPDH | This gene encodes a member of the glyceraldehyde-3-phosphate dehydrogenase protein family. The encoded protein has been identified as a moonlighting protein based on its ability to perform mechanistically distinct functions. The product of this gene catalyzes an important energy-yielding step in carbohydrate metabolism, the reversible oxidative phosphorylation of glyceraldehyde-3-phosphate in the presence of inorganic phosphate and nicotinamide adenine dinucleotide (NAD). The encoded protein has additionally been identified to have uracil DNA glycosylase activity in the nucleus. Also, this protein contains a peptide that has antimicrobial activity against *E. coli*, *P. aeruginosa*, and *C. albicans*. Studies of a similar protein in mouse have assigned a variety of additional functions including nitrosylation of nuclear proteins, the regulation of mRNA stability, and acting as a transferrin receptor on the cell surface of macrophage. Many pseudogenes similar to this locus are present in the human genome. Alternative splicing results in multiple transcript variants. | https://www.ncbi.nlm.nih.gov/gene/2597 |
| RPS13 | This gene encodes a ribosomal protein that is a component of the 40S subunit. The protein belongs to the S15P family of ribosomal proteins. It is located in the cytoplasm. The protein has been shown to bind to the 5.8S rRNA in rat. The gene product of the *E. coli* ortholog (ribosomal protein S15) functions at early steps in ribosome assembly. This gene is co-transcribed with two U14 small nucleolar RNA genes, which are located in its third and fifth introns. As is typical for genes encoding ribosomal proteins, there are multiple processed pseudogenes of this gene dispersed through the genome. | https://www.ncbi.nlm.nih.gov/gene/6207 |
| PSMC5 | PSMC5 (Proteasome 26S Subunit, ATPase 5) is a Protein Coding gene.the 26S proteasome is a multicatalytic proteinase complex with a highly ordered structure composed of 2 complexes, a 20S core and a 19S regulator. The 20S core is composed of 4 rings of 28 non-identical subunits; 2 rings are composed of 7 alpha subunits and 2 rings are composed of 7 beta subunits. The 19S regulator is composed of a base, which contains 6 ATPase subunits and 2 non-ATPase subunits, and a lid, which contains up to 10 non-ATPase subunits. Proteasomes are distributed throughout eukaryotic cells at a high concentration and cleave peptides in an ATP/ubiquitin-dependent process in a non-lysosomal pathway. An essential function of a modified proteasome, the immunoproteasome, is the processing of class I MHC peptides. This gene encodes one of the ATPase subunits, a member of the triple-A family of ATPases which have a chaperone-like activity. In addition to participation in proteasome functions, this subunit may participate in transcriptional regulation since it has been shown to interact with the thyroid hormone receptor and retinoid X receptor-alpha. Two transcript variants encoding different isoforms have been found for this gene. | https://www.ncbi.nlm.nih.gov/gene/5705 |
| MTCH1 | Mitochondrial Carrier 1; This gene encodes a member of the mitochondrial carrier family. The encoded protein is localized to the mitochondrion inner membrane and induces apoptosis independent of the proapoptotic proteins Bax and Bak. Pseudogenes on chromosomes 6 and 11 have been identified for this gene. Alternatively spliced transcript variants encoding multiple isoforms have been observed. Potential mitochondrial transporter. May play a role in apoptosis. | https://www.ncbi.nlm.nih.gov/gene/23787 |
| MKRN1 | Makorin Ring Finger Protein 1; This gene encodes a protein that belongs to a novel class of zinc finger proteins. The encoded protein functions as a transcriptional co-regulator, and as an E3 ubiquitin ligase that promotes the ubiquitination and proteasomal degradation of target proteins. The protein encoded by this gene is thought to regulate RNA polymerase II-catalyzed transcription. Substrates for this protein's E3 ubiquitin ligase activity include the capsid protein of the West Nile virus and the catalytic subunit of the telomerase ribonucleoprotein. This protein controls cell cycle arrest and apoptosis by regulating p21, a cell cycle regulator, and the tumor suppressor protein p53. Pseudogenes of this gene are present on chromosomes 1, 3, 9, 12 and 20, and on the X chromosome. Alternative splicing results in multiple transcript variants encoding different isoforms. E3 ubiquitin ligase catalyzing the covalent attachment of ubiquitin moieties onto substrate proteins. These substrates include FILIP1, p53/TP53, CDKN1A and TERT. Keeps cells alive by suppressing p53/TP53 under normal conditions, but stimulates apoptosis by repressing CDKN1A under stress conditions. Acts as a negative regulator of telomerase. Has negative and positive effects on RNA polymerase II-dependent transcription. | https://www.ncbi.nlm.nih.gov/gene/23608 |
| ERGIC3 | Possible role in transport between endoplasmic reticulum and Golgi. NOT MUCH KNOWN...RGIC3, which is regulated by miR-203a, is a potential biomarker for non-small cell lung cancer.https://www.ncbi.nlm.nih.gov/pmc/articles/PMC4638005/. ERGIC3 (Erv46 and ERp43) is located in the cis face of the Golgi apparatus and vesicular tubular structures between the transitional endoplasmic reticulum (ER) and cis-Golgi.10 ERGIC3 significantly affects cell growth and causes ER stress-induced cell death, and is involved in the invasion and metastasis in hepatocellular carcinomas (HCC).11,1 | https://www.ncbi.nlm.nih.gov/gene/51614 |
| ADRM1 | Adhesion Regulating Molecule 1; This gene encodes a member of the adhesion regulating molecule 1 protein family. The encoded protein is a component of the proteasome where it acts as a ubiquitin receptor and recruits the deubiquitinating enzyme, ubiquitin carboxyl-terminal hydrolase L5. Increased levels of the encoded protein are associated with increased cell adhesion, which is likely an indirect effect of this intracellular protein. Dysregulation of this gene has been implicated in carcinogenesis. Alternative splicing results in multiple transcript variants. [provided by RefSeq, Jul 2013]; Functions as a proteasomal ubiquitin receptor. Recruits the deubiquitinating enzyme UCHL5 at the 26S proteasome and promotes its activity. | https://www.ncbi.nlm.nih.gov/gene/11047 |
| TSPAN31 | The protein encoded by this gene is a member of the transmembrane 4 superfamily, also known as the tetraspanin family. Most of these members are cell-surface proteins that are characterized by the presence of four hydrophobic domains. The proteins mediate signal transduction events that play a role in the regulation of cell development, activation, growth and motility. This encoded protein is thought to be involved in growth-related cellular processes. This gene is associated with tumorigenesis and osteosarcoma. | https://www.ncbi.nlm.nih.gov/gene/6302 |
| DARS | Aspartyl-TRNA Synthetase; This gene encodes a member of a multienzyme complex that functions in mediating the attachment of amino acids to their cognate tRNAs. The encoded protein ligates L-aspartate to tRNA(Asp). Mutations in this gene have been found in patients showing hypomyelination with brainstem and spinal cord involvement and leg spasticity. Alternative splicing results in multiple transcript variants. Catalyzes the specific attachment of an amino acid to its cognate tRNA in a 2 step reaction: the amino acid (AA) is first activated by ATP to form AA-AMP and then transferred to the acceptor end of the tRNA. | https://www.ncbi.nlm.nih.gov/gene/1615 |
| RPL5 | Ribosomal Protein L5; The protein binds 5S rRNA to form a stable complex called the 5S ribonucleoprotein particle (RNP), which is necessary for the transport of nonribosome-associated cytoplasmic 5S rRNA to the nucleolus for assembly into ribosomes. The protein interacts specifically with the beta subunit of casein kinase II. Variable expression of this gene in colorectal cancers compared to adjacent normal tissues has been observed, although no correlation between the level of expression and the severity of the disease has been found. This gene is co-transcribed with the small nucleolar RNA gene U21, which is located in its fifth intron. As is typical for genes encoding ribosomal proteins, there are multiple processed pseudogenes of this gene dispersed through the genome. Required for rRNA maturation and formation of the 60S ribosomal subunits. This protein binds 5S RNA | https://www.ncbi.nlm.nih.gov/gene/6125 |
| RPL37A | Ribosomal Protein L37a; This gene encodes a ribosomal protein that is a component of the 60S subunit. The protein belongs to the L37AE family of ribosomal proteins. It is located in the cytoplasm. The protein contains a C4-type zinc finger-like domain. As is typical for genes encoding ribosomal proteins, there are multiple processed pseudogenes of this gene dispersed through the genome. | https://www.ncbi.nlm.nih.gov/gene/6168 |

## References

Eisenberg E, Levanon EY. Human housekeeping genes, revisited. Trends in Genetics 2013;29(10):569–74.

Janssens N, Janicot M, Perera T, et al. Housekeeping genes as internal standards in cancer research. Molecular Diagnosis 2004;8(2):107–13.

Ferlay J, Shin HR, Bray F, et al. Estimates of worldwide burden of cancer in 2008: GLOBOCAN 2008. International journal of cancer Journal international du cancer 2010;127(12):2893–917. doi:10.1002/ijc.25516.

Lallemant B, Evrard A, Combescure C, et al. Reference gene selection for head and neck squamous cell carcinoma gene expression studies. BMC Molecular Biology 2009;10(1):78.

De Jonge HJ, Fehrmann RS, de Bont ES, et al. Evidence based selection of housekeeping genes. PloS one 2007;2(9):e898.

Greer S, Honeywell R, Geletu M, et al. Housekeeping genes; expression levels may change with density of cultured cells. Journal of immunological methods 2010;355(1):76–79.

Du P, Kibbe WA, Lin SM. lumi: a pipeline for processing Illumina microarray. Bioinformatics 2008;24(13):1547–48.

Johnson WE, Li C, Rabinovic A., Adjusting batch effects in microarray expression data using empirical Bayes methods. Biostatistics 2007;8(1):118–27.

Krishnan N, Gupta S, Palve V, et al. Integrated analysis of oral tongue squamous cell carcinoma identifies key variants and pathways linked to risk habits, HPV, clinical parameters and tumor recurrence. F1000Res 2015;4:1215. doi: 10.12688/f1000research.7302.1

Krishnan NM, Dhas K, Nair J, et al. A Minimal DNA Methylation Signature in Oral Tongue Squamous Cell Carcinoma Links Altered Methylation with Tumor Attributes. Mol Cancer Res 2016;14(9):805–19. doi: 10.1158/1541-7786.MCR-15-0395.

Cancer Genome Atlas N. Comprehensive genomic characterization of head and neck squamous cell carcinomas. Nature 2015;517(7536):576–82. doi: 10.1038/nature14129.

Campos MS, Rodini CO, Pinto-Junior DS, et al. GAPD and tubulin are suitable internal controls for qPCR analysis of oral squamous cell carcinoma cell lines. Oral Oncol 2009;45(2):121–6. doi: 10.1016/j.oraloncology.2008.03.019.

Vandesompele J, De Preter K, Pattyn F, et al. Accurate normalization of real-time quantitative RT-PCR data by geometric averaging of multiple internal control genes. Genome biology 2002;3(7):research0034. 1.

Andersen CL, Jensen JL, Ørntoft TF. Normalization of real-time quantitative reverse transcription-PCR data: a model-based variance estimation approach to identify genes suited for normalization, applied to bladder and colon cancer data sets. Cancer research 2004;64(15):5245–50.

Pfaffl MW, Tichopad A, Prgomet C, et al. Determination of stable housekeeping genes, differentially regulated target genes and sample integrity: BestKeeper– Excel-based tool using pair-wise correlations. Biotechnology letters 2004;26(6):509–15.

Schmittgen TD, Livak KJ., Analyzing real-time PCR data by the comparative CT method. Nature protocols 2008;3(6):1101–08.

Xie F, Xiao P, Chen D, et al. miRDeepFinder: a miRNA analysis tool for deep sequencing of plant small RNAs. Plant molecular biology 2012;80(1):75–84.

Bustin SA, Benes V, Garson JA, et al. The MIQE guidelines: minimum information for publication of quantitative real-time PCR experiments. Clinical chemistry 2009;55(4):611–22.

Glare E, Divjak M, Bailey M, et al. β-Actin and GAPDH housekeeping gene expression in asthmatic airways is variable and not suitable for normalising mRNA levels. Thorax 2002;57(9):765–70.

Oliveira JG, Prados RZ, Guedes ACM, et al. The housekeeping gene glyceraldehyde-3-phosphate dehydrogenase is inappropriate as internal control in comparative studies between skin tissue and cultured skin fibroblasts using Northern blot analysis. Archives of Dermatological Research 1999;291(12):659–61.

Selvey S, Thompson EW, Matthaei K, et al. β-Actin—an unsuitable internal control for RT-PCR. Molecular and cellular probes 2001;15(5):307–11.

Zhong H, Simons JW., Direct comparison of GAPDH, β-actin, cyclophilin, and 28S rRNA as internal standards for quantifying RNA levels under hypoxia. Biochemical and biophysical research communications 1999;259(3):523–26.

Popovici V, Goldstein DR, Antonov J, et al. Selecting control genes for RT-QPCR using public microarray data. BMC bioinformatics 2009;10(1):42.

